# ELeFHAnt: A supervised machine learning approach for label harmonization and annotation of single cell RNA-seq data

**DOI:** 10.1101/2021.09.07.459342

**Authors:** Konrad Thorner, Aaron M. Zorn, Praneet Chaturvedi

## Abstract

Annotation of single cells has become an important step in the single cell analysis framework. With advances in sequencing technology thousands to millions of cells can be processed to understand the intricacies of the biological system in question. Annotation through manual curation of markers based on a priori knowledge is cumbersome given this exponential growth. There are currently ~200 computational tools available to help researchers automatically annotate single cells using supervised/unsupervised machine learning, cell type markers, or tissue-based markers from bulk RNA-seq. But with the expansion of publicly available data there is also a need for a tool which can help integrate multiple references into a unified atlas and understand how annotations between datasets compare. Here we present ELeFHAnt: *Ensemble learning for harmonization and annotation of single cells*. ELeFHAnt is an easy-to-use R package that employs support vector machine and random forest algorithms together to perform three main functions: 1) *CelltypeAnnotation* 2) *LabelHarmonization* 3) *DeduceRelationship*. *CelltypeAnnotation* is a function to annotate cells in a query Seurat object using a reference Seurat object with annotated cell types. *LabelHarmonization* can be utilized to integrate multiple cell atlases (references) into a unified cellular atlas with harmonized cell types. Finally, *DeduceRelationship* is a function that compares cell types between two scRNA-seq datasets. ELeFHAnt can be accessed from GitHub at https://github.com/praneet1988/ELeFHAnt.

## 1 Introduction

Single cell sequencing has become an important method for understanding biological systems at an increasingly granular level^1,2,3^. For single cell RNA-seq (scRNA-seq) data specifically, one of the primary questions is how to determine cell identity from the transcriptome. It is common to visualize such data and find clusters of cells with similarity in gene expression, but assigning each cluster a cell type is a much more open-ended task. Taking advantage of publicly available, annotated datasets in combination with supervised learning is a powerful method for addressing this question. Ensemble Learning for Harmonization and Annotation of Single Cells (ELeFHAnt) is an R package that utilizes this approach, enabling users to annotate clusters of single cells, harmonize labels across datasets to generate a unified atlas, and infer relationships among cell types between two datasets. It provides users with the flexibility of choosing between random forest and SVM (Support Vector Machine) based classifiers or letting ELeFHAnt apply both in combination to make predictions.

As an alternative to manual annotation, there are many automatic cell annotation tools currently available based on either gene marker, correlation, or machine learning-based methods, each with varying levels of performance^4^. The label transfer method from Seurat is among the most well-known, quickly identifying labels by finding neighboring “anchor” cells^1^. There are also deep learning-based tools emerging such as scANVI, which utilizes generative models but requires greater computation time^5^.

ELeFHAnt is a supervised machine learning-based tool that enables researchers to identify cell types in their scRNA-seq data while providing additional unique features. ELeFHAnt gives users the ability to use and compare not just one but multiple classification algorithms simultaneously through its ensemble method, weighting their predictions to produce the best consensus among them. SVM and random forest classifiers were selected for the ensemble based on their superior accuracy and computation time in a benchmarking study^6^. Additionally, selecting the optimal reference is a challenge addressed by harmonization, that allows users to integrate multiple datasets together into an atlas. A standardized set of labels is generated across all of them, which can subsequently be used to annotate new datasets. Relationships between datasets can also be deduced to better understand how each was annotated. This is provided in an easily interpretable heatmap format that compares all cell types between them. Finally, a subsampling procedure is used to enable faster predictions while being shown not to influence reproducibility.

ELeFHAnt has been tested on multiple public datasets related to early fetal development and the human pancreas, where it was able to effectively predict and harmonize cell types. We then present a case study for when reference and query are not as directly related, specifically using a tissue atlas to annotate organoid data. We also perform benchmarking and find that our approach achieves comparable performance to other annotation tools. Taken together, we hope this demonstrates that ELeFHAnt lends itself to various problems in the single cell research space.

## 2 Results

### 2.1 Label harmonization creates a unified atlas from multiple datasets

When integrating single cell RNA datasets with different sets of cell type assignments/labels, it can become very difficult to infer the cell type label that an integrated cluster should be associated with. Many cell labels can be found to overlap or disagree with one other. To solve this problem, we designed a function (*LabelHarmonization*) to harmonize cell labels from multiple datasets into a unified atlas with cell labels assigned to each cluster of cells in the integrated dataset. To demonstrate ELeFHAnt’s *LabelHarmonization* we used three datasets with intestinal cells profiled from fetal gut development: E-MTAB-8901, GSE158702, and E-MTAB-10187 (Table 1). Briefly, we integrate the three atlases (~120k cells) using Seurat’s canonical correlation analysis-based integration algorithm, resulting in 41 clusters. For label harmonization we kept 120 of the 141 total cell labels (~112k cells), removing cell labels that were not informative (Supplementary Table 2), and then created training and test sets using 60% and 40% of the data (please see Methods for details). Using its ensemble learning method ELeFHAnt predicted cell labels for each of the 41 integration clusters, harmonizing the 120 cell labels to a final set of 33 (Figure 2). In Figure 2, the left panel shows integrated cells labeled with the 120 cell labels that all references contributed, whereas on the right panel, integrated cells labeled with the 33 harmonized cell labels generated by ELeFHAnt are shown. As an example, there are initially ten different neuronal cell labels as shown on the left, while on the right ELeFHAnt harmonized these cell labels and found three that best annotate the given clusters.

**Table 1:** Attributes of the public single cell RNA-seq datasets used in the analyses.

| Dataset | Organ | Number of cells | Number of features/genes | Number of clusters | Number of cell types | Sequencing protocol |
| --- | --- | --- | --- | --- | --- | --- |
| E-MTAB-8901<sup>7</sup> | Duodenum | 21592 | 33694 | 31 | 26 | 10X |
| GSE158702<sup>8</sup> | Intestine | 20779 | 33538 | 29 | 94 | 10X |
| E-MTAB-10187<sup>9</sup> | Intestine | 77866 | 28691 | 38 | 24 | 10X |
| E-MTAB-10187<sup>9</sup> | Esophagus, Lung, Liver, Stomach and Intestine | 155232 | 28691 | 42 | 27 | 10X |
| E-MTAB-10268<sup>9</sup> | Transplanted HIO | 1868 | 22016 | 16 | 15 | 10X |
| GSE84133<sup>10</sup> | Pancreas | 8569 | 20125 | 18 | 14 | inDrop |
| E-MTAB-5061<sup>11</sup> | Pancreas | 3514 | 25525 | 16 | 15 | Smart-Seq2 |
| GSE83139<sup>12</sup> | Pancreas | 635 | 19950 | 9 | 8 | SMARTer |
| GSE81608<sup>13</sup> | Pancreas | 1600 | 39851 | 11 | 8 | SMARTer |

**Figure 1:**
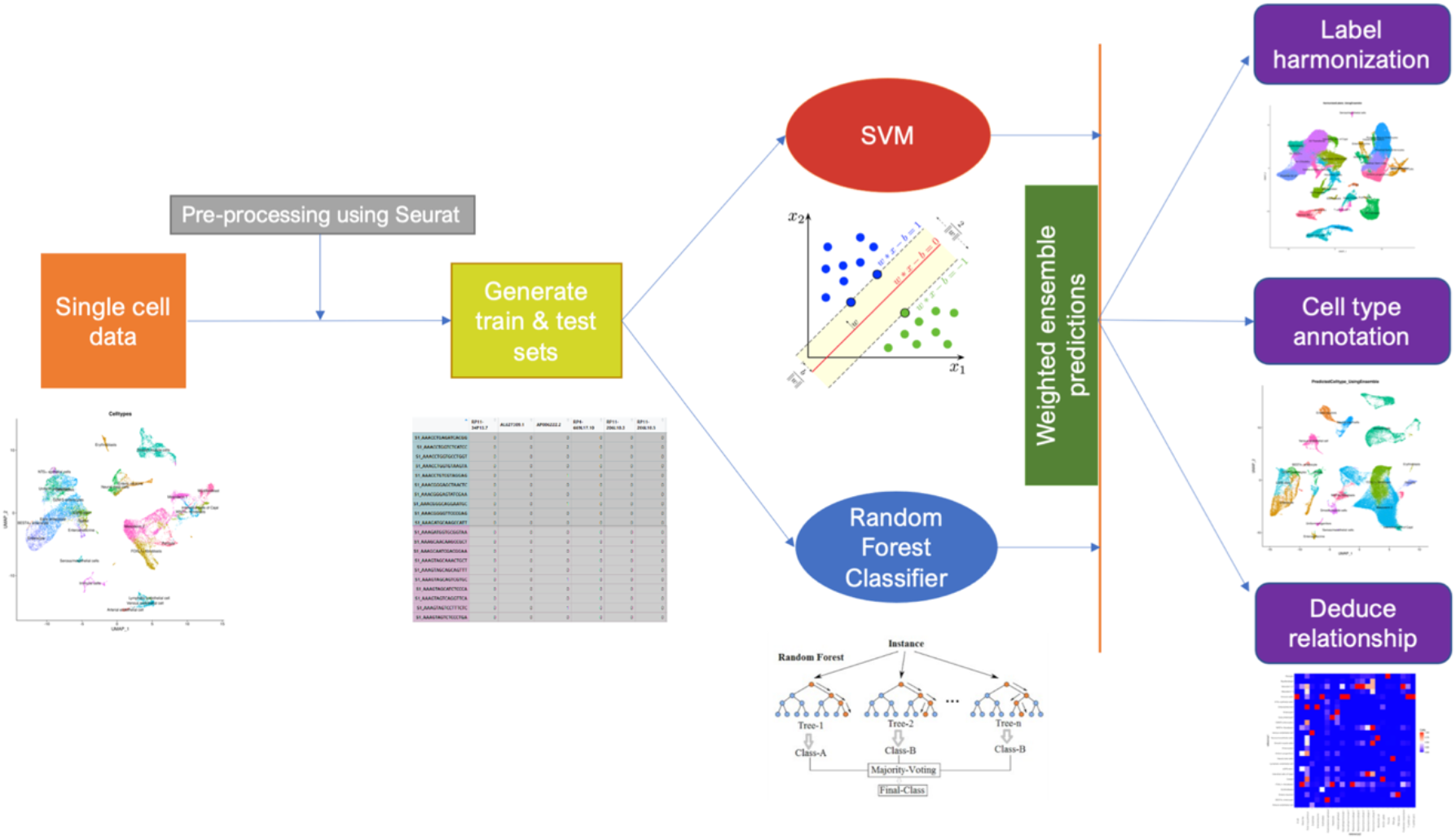
ELeFHAnt model: Workflow summary of different functions.

**Figure 2:**
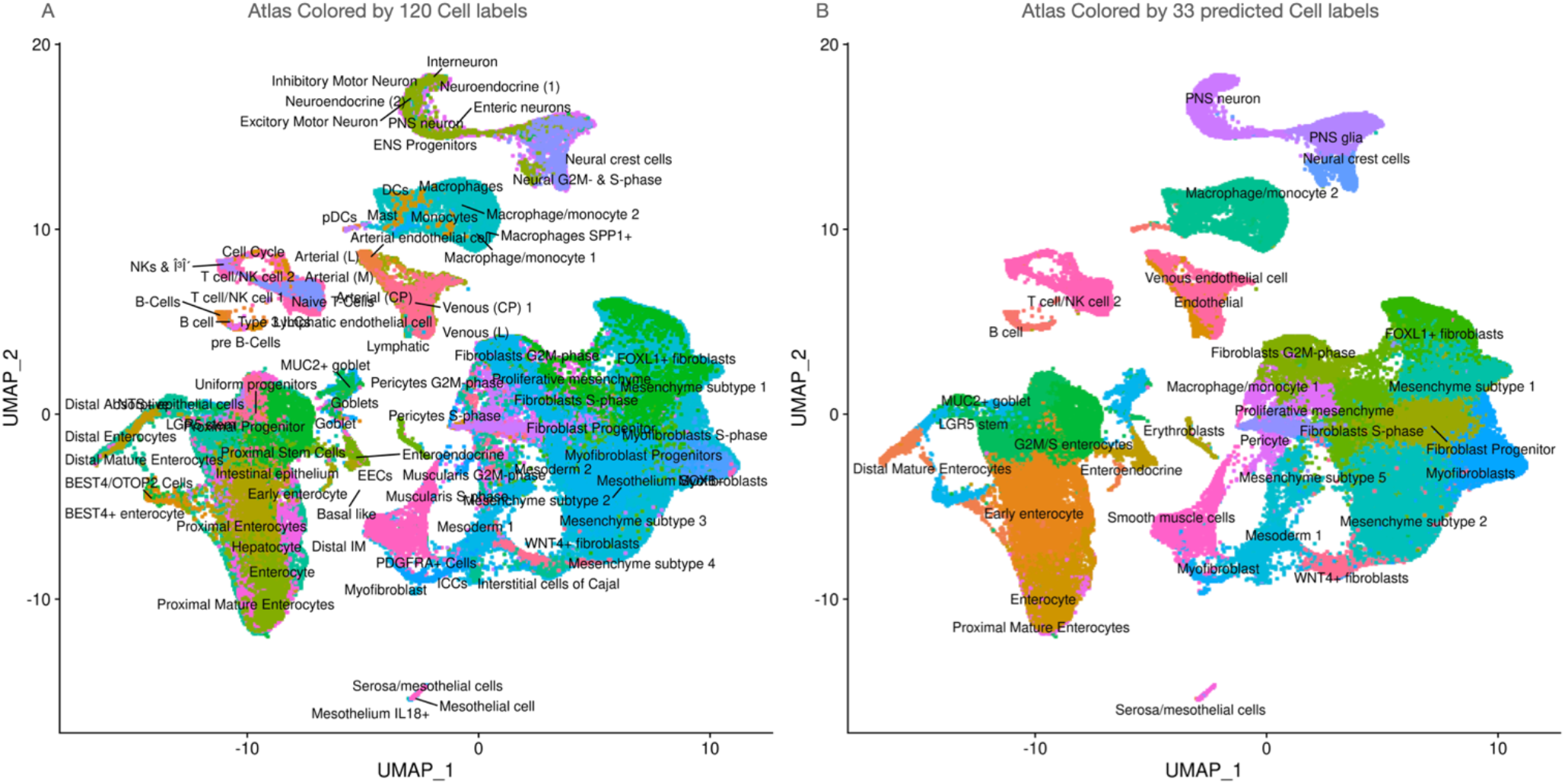
ELeFHAnt generates harmonized cell labels : ~112k cells are integrated from three fetal gut references. A) Atlas colored using 120 cell labels B) Atlas colored using 33 cell labels predicted by ELeFHAnt

Whether these harmonized cell labels are appropriate for each cluster can be assessed by the *validatePredictions* function in ELeFHAnt, which uses gene set enrichment analysis (GSEA) as its foundation. The top 100 markers from each cell label in the integrated dataset are used as gene sets and their enrichment is tested in the top 100 markers for each integrated cluster (please see Methods for details). This GSEA pre-ranked based enrichment analysis shows that ELeFHAnt’s predictions are among the top choices based on adjusted p-value and/or size of the gene set enriched in leading edge analysis (Supplementary Table 1, (Harmonized_Labels_GSEA_Stats)). We also provide complete enrichment results from GSEA (Supplementary Table 1, (GSEA Analysis Validate Pred.) and have highlighted predicted label choices from ELeFHAnt.

### 2.2 Deduce Relationship reveals similarity in cell types across datasets

With the growing number of scRNA-seq datasets available, hypothesis generation can be piloted by utilizing multiple reference scRNA datasets. Comparing reference datasets, or in-house datasets to publicly available datasets, is important for facilitating experimental design and assessing differences in annotation, sequencing, or clustering The *DeduceRelationship* function in ELeFHAnt aids in this process by comparing the cell types or other metadata between two datasets and calculating their relative similarity. To demonstrate, we compared E-MTAB-8901 (reference1) vs E-MTAB-10187 (intestine) (reference2), where we down sample both the datasets to 300 cells per cell type and used ELeFHAnt’s ensemble learning method (Figure 3). We can clearly see that “Mesoderm2” cell type in reference1 corresponds to many of the mesenchyme subtypes in reference2, and similarly all immune sub-cell types in reference2 are similar to “Immune cells” cell type in reference1.

**Figure 3:**
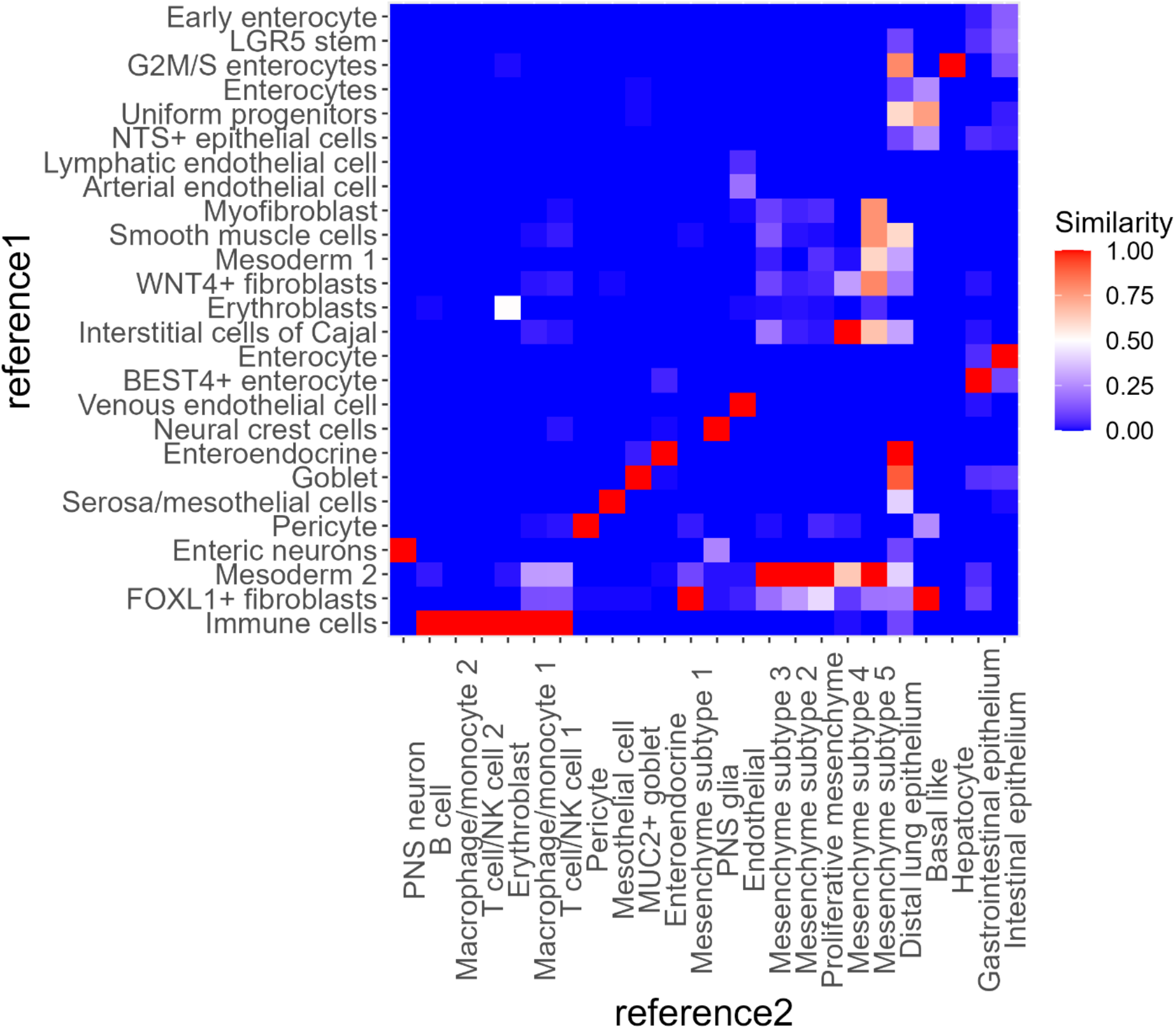
ELeHAnt infers relative similarity among cell types from two references :The output of Deduce Relationship represents the confusion matrix as a heatmap, with each square signifying how many cells of a given cell type in one reference were classified as a cell type in the other. It is normalized such that each cell type in reference 2 has a red square that shows the most closely related cell type in reference 1

### 2.3 Cell Type Annotation uses references to label unannotated cells

Cell type annotation in ELeFHAnt is performed by two sub functions: 1) *ClassifyCells* 2) *ClassifyCells_usingApproximation.* The first is annotation performed on a per-cell basis, whereas the second assigns each cluster in the query the most frequently predicted cell type for that cluster (see Methods for details). This per-cluster approximation generally has more distinct cell populations and is recommended when the reference is not ideal in terms of size or level of annotation.

To showcase ELeFHAnt’s cell type annotation function, we used two different scenarios 1) reference and query are downsampled to a similar of number of cells 2) reference and query are unaltered. For testing in first scenario, we down sampled dataset E-MTAB-8901 to 300 cells per cell type (~6500 cells) and set it as the reference (Figure 4A). We then downsampled E-MTAB-10187 to 200 cells per cluster (~7600 cells) and set it as query (Figure 4B). For second scenario, all cells in E-MTAB-8901(~21k cells) (Figure 4A) were set as reference and all cells in E-MTAB-10187 (~77k cells) (Figure 4B) were set as query. For both scenarios, we used ELeFHAnt’s ensemble learning with the *ClassifyCells* function to predict cell types for each cell (Figure 4D and Figure 4E). Scenario 2 clearly exhibits some misclassification compared to scenario 1 (Figure 4D). Cluster 35 in the query was labelled as “Erythroblasts” and “Immune cells” by ELeFHAnt in scenario 1, matching the known cell types in E-MTAB-10187 (intestine). However, in scenario 2 “Erythroblasts” are also present in a distant cluster 10 that was labelled as “Mesoderm 1” in scenario 1.

**Figure 4:**
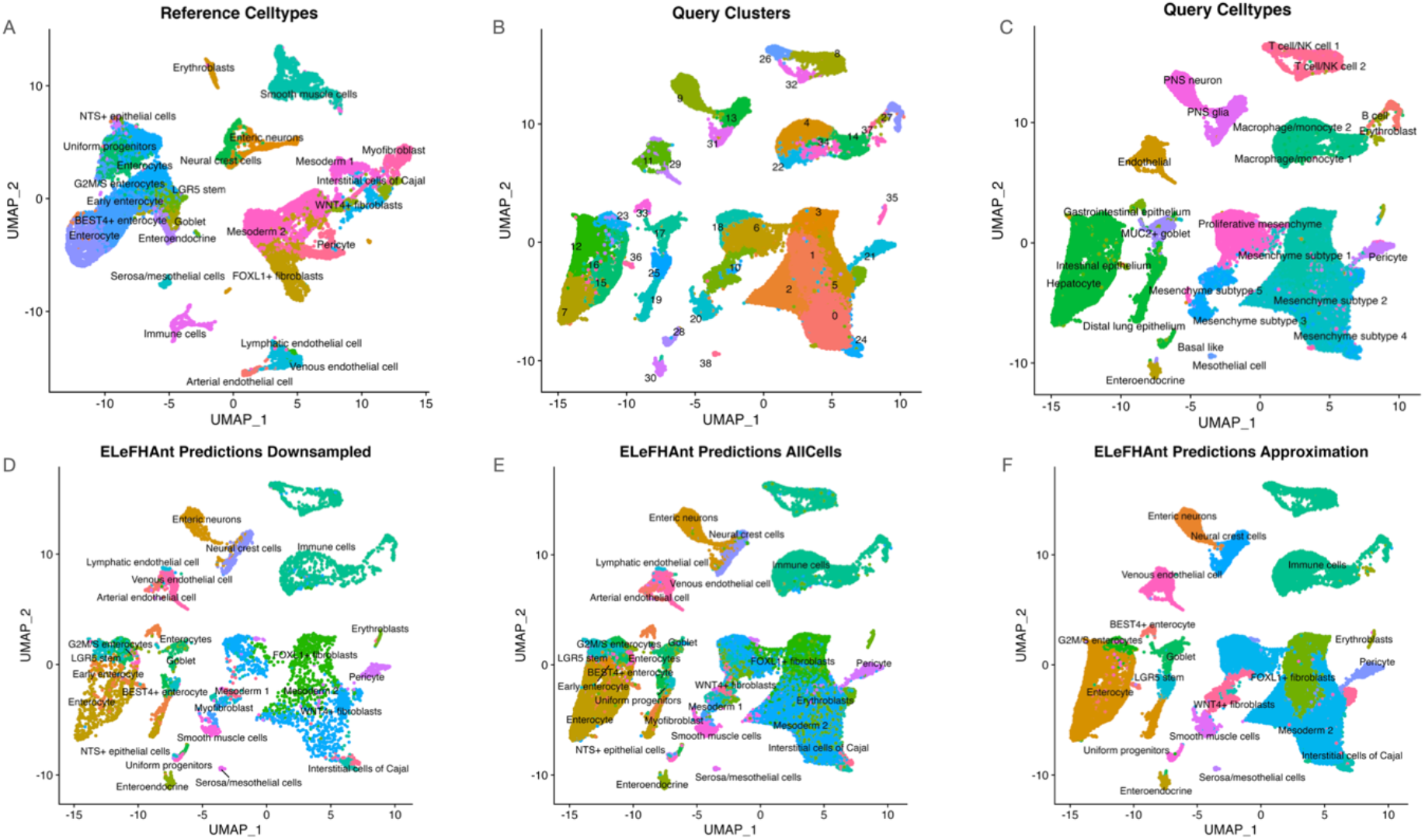
Cell type annotation using ELeFHAnt : ~21k cells in reference (4A) and ~77k cells in query (4B). 4D, 4E and 4F show the ELeFHAnt predictions for 1) down sampled reference and query 2) all cells in reference and query 3) approximation-based predictions for all cells in reference and query

This prompted us to the *ClassifyCells_usingApproximation* function where ELeFHAnt finds the best prediction for each query cluster rather than each cell (Figure 4F). ELeFHAnt annotated the 38 query clusters with a total of 19 reference cell types. Comparing the annotations in 4D and 4F to the known cell types in 4C, we see a 1:1 correspondence across broad cell types including endothelial, epithelial, mesenchymal, and neuronal. Although total number of cell types is reduced, from a biological standpoint the annotation is correct. We show all the choices available to ELeFHAnt using the number of cells shared among clusters and predicted cell types (Supplementary Figure 1). Please refer to benchmarking section to compare the cell type predictions on the same data using ELeFHAnt, scPred, Seurat label transfer, and scANVI.

To further demonstrate the cell type annotation functionality of ELeFHAnt we used the following human pancreatic datasets: GSE84133, E-MTAB-5061, GSE83139, and GSE81608 (Table 1). These datasets are regularly used for benchmarking integration, cell type annotation, and other techniques. GSE84133 was chosen as the reference to annotate cell types in the remaining datasets. We applied ELeFHAnt’s ensemble learning approach with *ClassifyCells* to train on the reference and predict cell types in each of the query datasets, and results are shown in Supplementary Figures 2 and 3.

Finally, we present a case study to show an application of ELeFHAnt in a possible experimental setup. E-MTAB-10187 (~155k cells) is used as the reference (Figure 5A), an atlas studying fetal development across 5 organs (Esophagus, Lung, Liver, Stomach and Intestine). E-MTAB-10268 (1868 cells), a dataset of in-vitro transplanted human intestinal organoids (tHIOs), is used as the query (Figure 5B). We first compare the two datasets using the *DeduceRelationship* function to infer relative similarity between the cell types among the two datasets (Figure 5C). We then learned the cell types from the reference and predicted the cell types for query, with ensemble learning and the *ClassifyCells* approach (Figure 5D). ELeFHAnt was able to annotate mesenchymal subtypes, epithelial, basal, ciliated, and endocrine cell types in the tHIO dataset (Figure 5).

**Figure 5:**
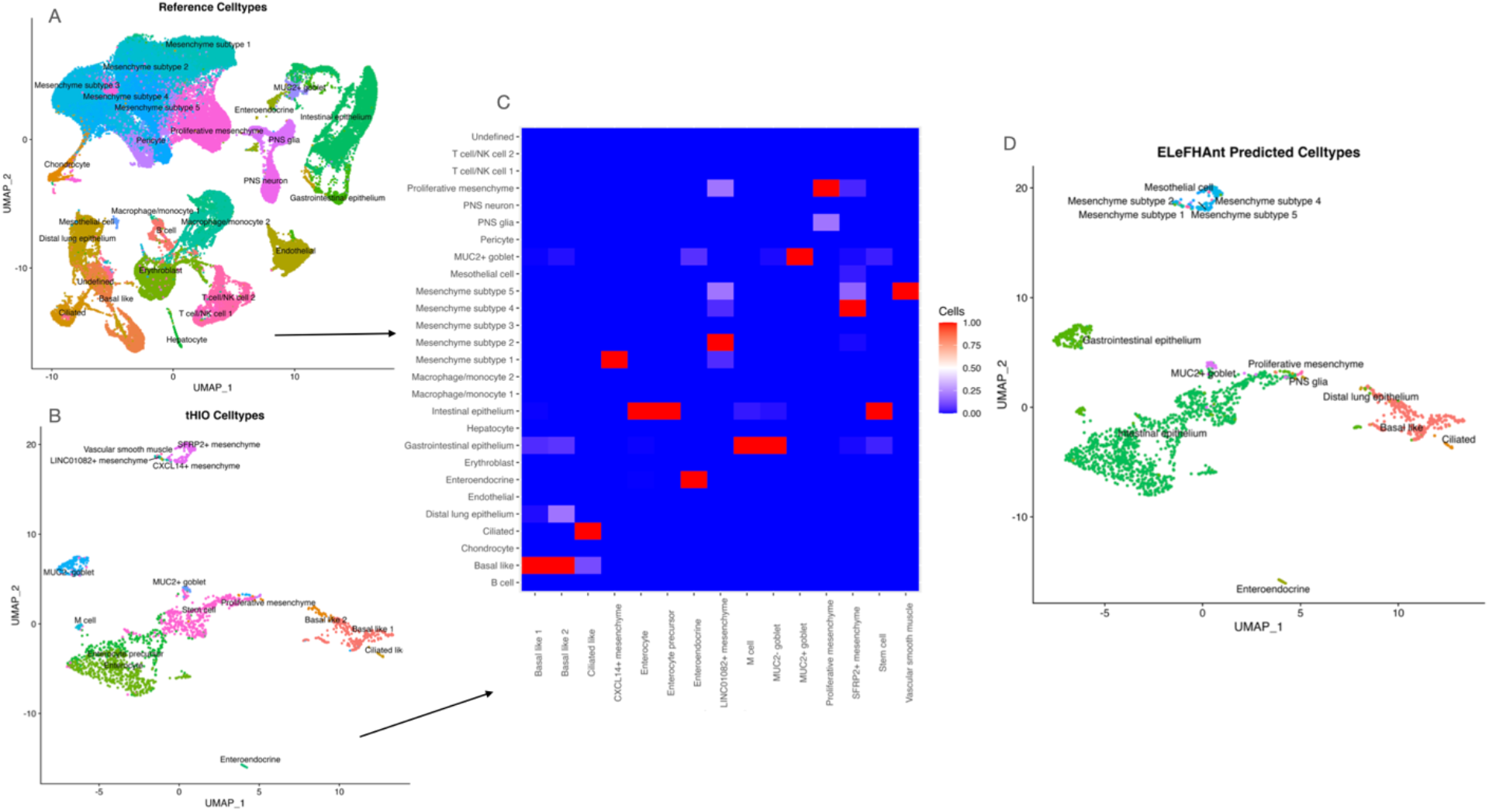
Case Study :A) ~155k cells with 27 cell types in reference B) 1868 cells with 15 cell types in query C) relative similarity based on number of cells shared among cell types in reference and query D) Cell type annotation based cell types learnt from reference

### 2.4 Subsampling has reproducible predictions and enables scalability

To assess reproducibility, each function was run 3 times with varying numbers of cells used during subsampling, at increments of 100, 300 and 500. This will also help determine what number of examples during training leads to the best performance for the SVM and random forest classifiers. E-MTAB-8901was used as the reference and E-MTAB-10187 (intestine) was used as the query (please see Table 1). Each pair of tests was compared to measure the percent of cells with same annotation, or in the case of *DeduceRelationship* the percent of cell types in one reference matched to the same cell type in the other reference. For annotation, we used *ClassifyCells_usingApproximation* approach and the predicted cell labels assigned to the clusters were >96% consistency (percent agreement) when subsampling 100, 300 and 500 cells (Supplementary Figure 4A). In the case of deduce relationship we also see little variability across tests, with similarity measurements remain approximately 95% consistent (Supplementary Figure 4B). However, the annotations determined by label harmonization were only 55-70% consistent ((Supplementary Figure 4C). Relative to the other two functions, the complexity from the number of datasets and cell types suggests that harmonization does benefit from a greater number of cells used (Supplementary Figure 2). This is seen in how agreement was highest when subsampling 300 and 500.

As an additional test, the functions were run three consecutive times with the same number of cells subsampled. As anticipated, there was 100% agreement, which is related to having constant random seeds that prevent variation during subsampling and training.

**Table 2:**
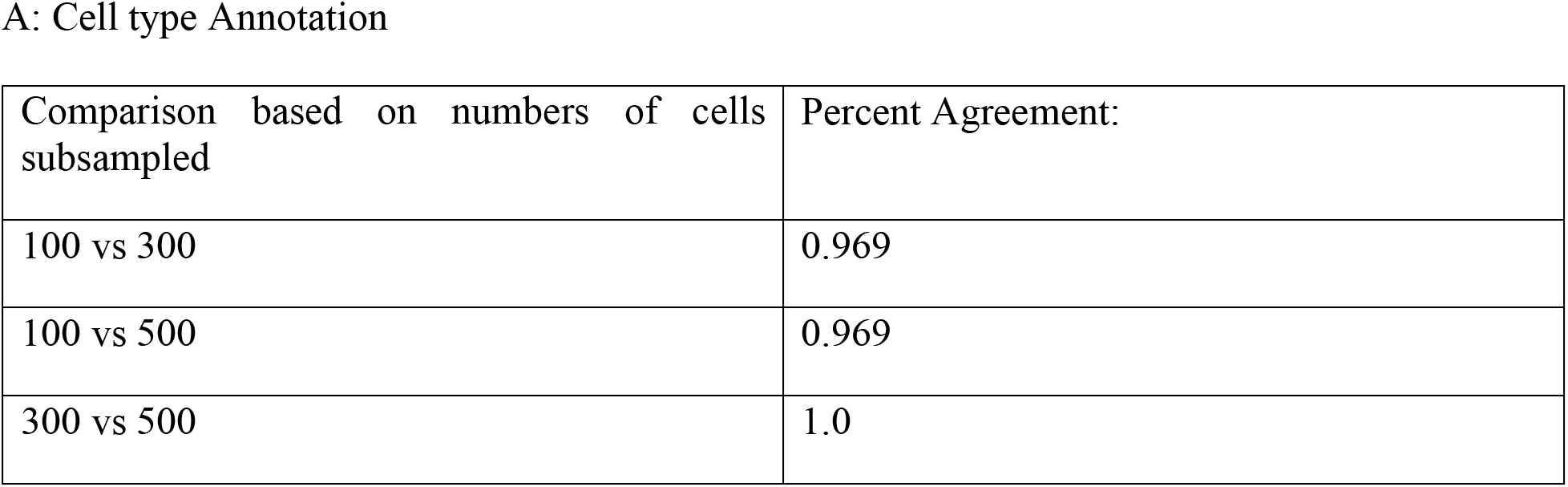
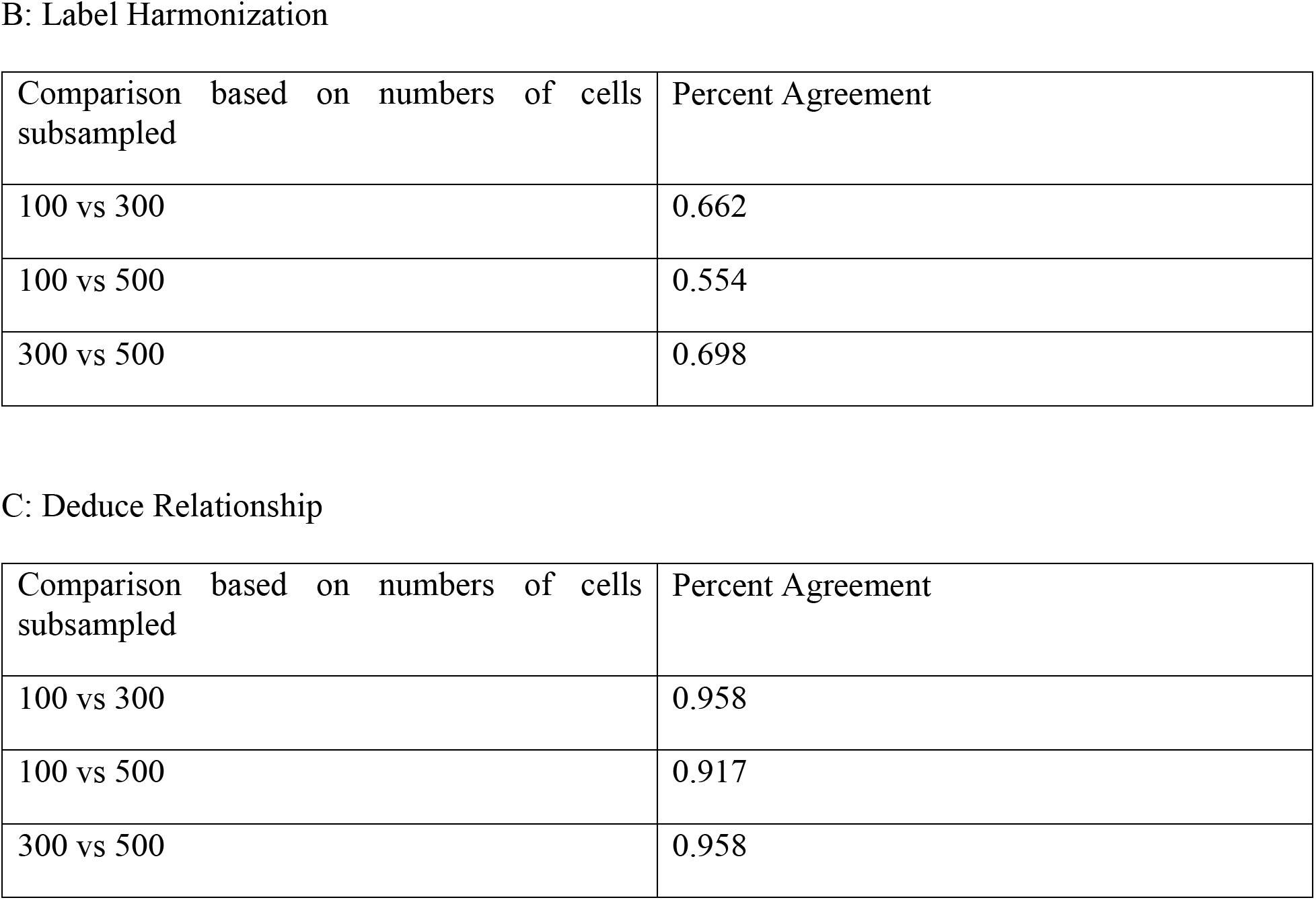
Comparing annotations between tests for (A) Annotation (B) Harmonization (C) Deduce Relationship.

### 2.5 ELeFHAnt’s performance in annotation is comparable to other tools

To measure the performance of ELeFHAnt, three tools were selected based on their performance relative to other tools in a benchmarking^6^, as well as their usage of different algorithms. Only cell annotation was assessed because this is the only functionality all four have in common. scPred is a cell annotation tool that utilizes SVM as its classifier, but departs from ELeFHAnt by using a radial kernel^14^. Additionally, the features used in training are principal components generated by single value decomposition (SVD) of the gene expression matrix. scANVI meanwhile is an extension of a previous tool called scVI designed for scRNA-seq integration, both of which employ probabilistic models^5^. It uses a form of deep learning with generative models, and requires that the reference and query be harmonized together prior to annotation. Finally, Seurat’s label transfer is based on identifying “anchors” between the reference and query in a low-dimensional space, an idea closely related to mutual nearest neighbors^1^. From these descriptions, ELeFHAnt’s cell annotation differs in multiple respects. The query and reference are neither processed separately nor integrated together, but rather they are merged prior to normalization and scaling. Also, ELeFHAnt does not use dimensionality reduction for its features, but uses the expression of genes identified as highly variable directly. Finally, ELeFHAnt can use up to two different classifiers when using its ensemble method.

The benchmarking was designed to test the annotation within and between datasets, with a focus on cell types in early human development. The reference used is E-MTAB-8901 while the query used is E-MTAB-10187 (intestine). It is important to note that the query has nearly three times the number of cells as the reference (77,672 versus 21,592), but they maintain a comparable number of cell types (26 versus 24). For the inter-dataset benchmark, the reference was downsampled to 300 cells per cell type and the query to 200 cells per cluster, and per-cell annotation with *ClassifyCells* was performed (Figure4A-4D). For the intra-dataset test, only the downsampled reference was used, but it was divided into training and test sets composed of 70 and 30 percent of the data (Supplementary Figure 5). scANVI results are provided in supplementary figures (Supplementary Figure 7).

#### 2.5.1 Intra-dataset

The intra-dataset test is more straightforward given that there are no batch effects or differences in annotation between datasets. The benchmarking showed ELeFHAnt, scPred, and Seurat perform equally well, with approximately 90% correspondence to the known cell types. ELeFHAnt had the highest overall accuracy at 90.5% while scPred had the lowest at 87.5% but a higher precision. This is related to scPred’s rejection option that led to a small population being labeled as “unassigned”, where the other tools had multiple cell types seen to be close together (Supplementary Figure 8). The predictions of scANVI diverged from the other tools, as only 20 out of the 26 known cell types were identified. In most cases these omissions were explained by labelling closely related cells together, such as labelling smooth muscle cells as myofibroblasts. Especially for the cluster of epithelial cells, scANVI struggled to distinguish between cell types and omitted many of interest, such as goblet cells.

**Table 3:** Intra-dataset accuracy, precision, and recall for three annotation tools. Precision and recall values are weighted macro-averages across cell types.

| Annotation Tool | Accuracy | Precision | Recall |
| --- | --- | --- | --- |
| ELeFHAnt | 0.905 | 0.906 | 0.905 |
| scPred | 0.875 | 0.941 | 0.875 |
| Seurat label transfer | 0.899 | 0.903 | 0.899 |

#### 2.5.2 Inter-dataset

The inter-dataset test meanwhile shows overall similar results between ELeFHAnt, scPred, and Seurat. Performance was measured by comparing the individual cell annotations of each tool in a pairwise fashion. ELeFHAnt and Seurat are shown to compare favorably to one another with 89.6% agreement. As with the intra-dataset test, scPred left some cells as “unassigned”, which may be beneficial if not all cell types in the query are represented in the reference (Supplementary Figure 6). At the same time, scPred was least consistent with the other tools, as seen with NTS+ epithelial cells and BEST4+ enterocytes, which may indicate misclassifications. scANVI cannot be compared to the other three tools as directly due to differences in the UMAP, but it also captured many of the same cell types. The largest difference is that all endothelial cells were labeled as venous endothelial, whereas the other tools identified all three subtypes.

Overall, the results of these benchmarking tests show that ELeFHAnt and Seurat’s label transfer are best in terms of accuracy and efficiency for identifying cell types. Additionally, despite downsampling the data ELeFHAnt can achieve comparable results to the other tools.

**Table 4:** Fraction of cells with same annotation between each pair of tools during inter-dataset benchmarking. Reference and query were downsampled, and per-cell annotation was performed.

| Annotation Tool Comparison | Fraction of annotations in common |
| --- | --- |
| ELeFHAnt vs scPred | 0.792 |
| ELeFHAnt vs Seurat label transfer | 0.896 |
| scPred vs Seurat label transfer | 0.801 |

## 3 Discussion

We have introduced the R package ELeFHAnt, a supervised learning approach for finding cell types in single cell data. ELeFHAnt should be familiar to users of Seurat, and requires only single line commands to use, making it approachable for researchers. It implements an ensemble approach to cell annotation not found in other widely used tools, while also bringing additional functionalities to the single cell research space. For annotation, we recognize that as with any machine learning task the training data is as important as the algorithm itself, and recommend that users take the time to compare datasets and consider harmonizing them. To do so, *DeduceRelationship* can be used to compare cell types or other metadata between two datasets, while *LabelHarmonization* allows users to take advantage of the information in multiple reference datasets when there is no “best” choice. These methods could prove beneficial even when used in conjunction with other annotation tools. ELeFHAnt also has the flexibility of using two different classifiers with the ability to combine their results. Users will benefit from seeing how SVM, random forest, and ensemble predictions compare for their datasets as they can capture different aspects of the data. Despite this added complexity, we showed that when downsampling to 500 cells or less per cell type we can achieve results on par with other tools that utilize entire datasets. Even in situations where there is a less than optimal reference, ELeFHAnt uses a unique approximation approach to label clusters in the query.

One potential area of improvement would be the validation approach. In the label harmonization example, there were two instances where GSEA showed no enrichment for ELeFHAnt’s predicted cell types: cluster 1, labeled as fibroblast progenitor, and cluster 39, labeled as macrophage/monocyte 1 (Supplementary Table 1). By comparing the ensemble learning confusion matrix produced by ELeFHAnt to the GSEA results, we noted that mesenchyme subtype 5 (15th choice in ELeFHAnt) was enriched for cluster 1 and none of the ELeFHAnt choices were enriched for cluster 39. This could potentially be addressed by using expression-based enrichment tests instead of the current pre-ranked gene-based enrichment.

There are plans for a Shiny version of ELeFHAnt, providing a simple user interface for using these functions, as well as a Python version that works within a Scanpy-based framework. The approach used by ELeFHAnt could also be applied to other analyses of single cell data. For example, the same machine learning concepts can be used for pseudo time analysis, harnessing reference datasets to understand cell states and the transitions between them. Also, with Seurat’s ability to handle multi-modal data, these methods could be used with scATAC-seq and other epigenetic datasets. Finally, we anticipate that these methods could also be applied to the deconvolution of bulk RNA-seq, learning from scRNA-seq datasets to identify their cell type composition.

## 4 Methods

### 4.1 Data pre-processing

All datasets must begin as or be converted to the *SeuratObject* format, and specifically for the reference datasets contain a column in the metadata named “Celltypes”. If *downsample* is set equal to “TRUE”, each dataset is subset to the value specified by *downsample_to* for every cell type or cluster. The commands *NormalizeData*, *FindVariableFeatures*, and *ScaleData* are then applied. *NormalizeData* applies a log-normalization that corrects for differences in read depth, *FindVariableFeatures* selects genes with high cell-to-cell variation that reflect more biologically meaningful genes (can be changed using *selectvarfeatures*), and *ScaleData* scales and centers the expression values.

### 4.2 SVM

The SVM algorithm was implemented using the R package *e1071*^15^. The goal of SVM is to find within n-dimensional space hyper-planes that separate the data into two classes. The “support vectors” are the data points closest to hyper-planes, with the distance between them defining a margin, and the optimal hyper-planes are those which maximize this margin. For our implementation, it is assumed the data is linearly separable and therefore a linear kernel is used. By default, the regularization or “cost” is set to 10 that controls the degree of misclassification tolerated to prevent overfitting, and a 10-fold cross validation is performed to measure accuracy of the model. Cell type annotation typically involves multiple cell types, in which case the algorithm uses multiple binary classifiers in a “one-versus-one” approach for classification.

### 4.3 Random forest

The random forest algorithm was implemented using the R package *randomForest*^16^. It consists of an ensemble of decision trees (by default 500), a decision tree being a model that classifies data by learning rules about its features. Each tree differs by randomly sampling from the training dataset with replacement, and similarly for every split in every tree only a random sample of the features is considered. During testing each tree will return its own prediction, and the final output is determined by majority vote among the trees.

### 4.4 Weighted ensemble

For *LabelHarmonization*, *ClassifyCells_usingApproximation*, and *DeduceRelationship*, the predictions generated by the SVM and random forest classifiers are represented as two confusion matrices, with reference cell types on one axis and query clusters on the other. Both classifiers also have an accuracy calculated, for SVM determined during cross-validation on the training data, and for random forest from the out of box error rate. When *classification.method* is set to “ensemble”, each confusion matrix is multiplied by its accuracy value as a “weight” on its predictions. The SVM and random forest matrices are then normalized by dividing each by their max value. Finally, the consensus confusion matrix is the sum of the two matrices. *ClassifyCells* differs in that it generates a table with each cell in the test set having a predicted cell type. In this scenario, the predictions of SVM and random forest are compared, and if they are not equal the cell type from the classifier with greater accuracy is kept.

### 4.5 Label Harmonization

Harmonization requires either an integrated *SeuratObject*, or a minimum of two datasets that are to be combined using Seurat’s canonical correlation analysis (CCA) based integration (Stuart et al., 2019). Briefly, CCA performs dimensionality reduction, L2 normalization is applied to the canonical correlation vectors, and mutual nearest neighbors (MNNs) are identified. These MNNs, or integration “anchors” between datasets, are used to correct for batch effects. The combined dataset then undergoes scaling, dimensionality reduction, and clustering. For the harmonization itself, stratified train and test sets are generated using 60 and 40 percent of the data, respectively. Finally, the specified *classification.method* is run.

### 4.6 Cell type Annotation

Cell annotation consists of two methods: *ClassifyCells_usingApproximation* and *ClassifyCells*. For both methods, the query and reference are merged and undergo pre-processing. For *ClassifyCells_usingApproximation* only, predictions are converted into a confusion matrix with cell types from the reference on one axis and clusters from the query on the other. The confusion matrix is normalized by row such that the max value for each cluster determines the best corresponding cell type. A copy of the confusion matrix is saved to a separate text file. For *ClassifyCells*, the predictions for each cell are left unmodified. The final output is the query with predicted cell types added to the metadata.

### 4.7 Deduce Relationship

As the first step, *DeduceRelationship* takes two datasets referred to as reference1 and reference2 and merges them into a single dataset. Following pre-processing, the training set is generated from reference1 and the test dataset from reference2, and the specified *classification.method* is run. The resulting confusion matrix is normalized per row such that each cell type in reference2 has a maximum value in reference1 that represents the best corresponding cell type. Finally, the confusion matrix is displayed as a heatmap and saved.

### 4.8 Validation of Predictions

The output of *CelltypeAnnotation* and *LabelHarmonization*can can be checked using *ValidatePredictions*, which is run automatically when *validatePredictions* is “TRUE”. For the reference or integrated atlas, it finds differential markers for each cell type with Seurat’s *FindAllMarkers* method, which uses a Wilcox rank sum test by default. The top 100 markers for each cell type are selected and split into individual gene sets. Next, for the query or integrated atlas it finds the top 100 markers for each Seurat cluster and ranks them based on average Log2 fold change. Gene set enrichment analysis (GSEA) with the PreRanked method is used to calculate statistical enrichment of gene sets in either end of the ranking in the query or integrated Seurat cluster ranked markers. The final GSEA results are saved to a text file.

### ELeFHAnt Reference Plugins

We provide 10 pre-processed reference datasets for users to get started with utilizing ELeFHAnt functionalities. Reference plugins can be accessed using: https://www.dropbox.com/sh/6hd2skriqqlokwp/AAAVol-_qPlCdA4DpERWjkeJa?dl=0. We plan to add more processed datasets in the future.

## Code Availability

All the source code, tutorials and version updates can be accessed through GitHub: https://github.com/praneet1988/ELeFHAnt

## Acknowledgements

We would like to thank Drs. Anil Jegga, Emily Miraldi, and Nathan Salomonis at Cincinnati Children’s Hospital Medical Center for their valuable feedback.

## Author Contributions

K.T. and P.C. designed, implemented, tested the algorithm, and wrote the manuscript. A.M.Z. provided feedback for testing and improving the algorithm from a research perspective.

## Supplementary Figures

**Supplementary Figure1:**
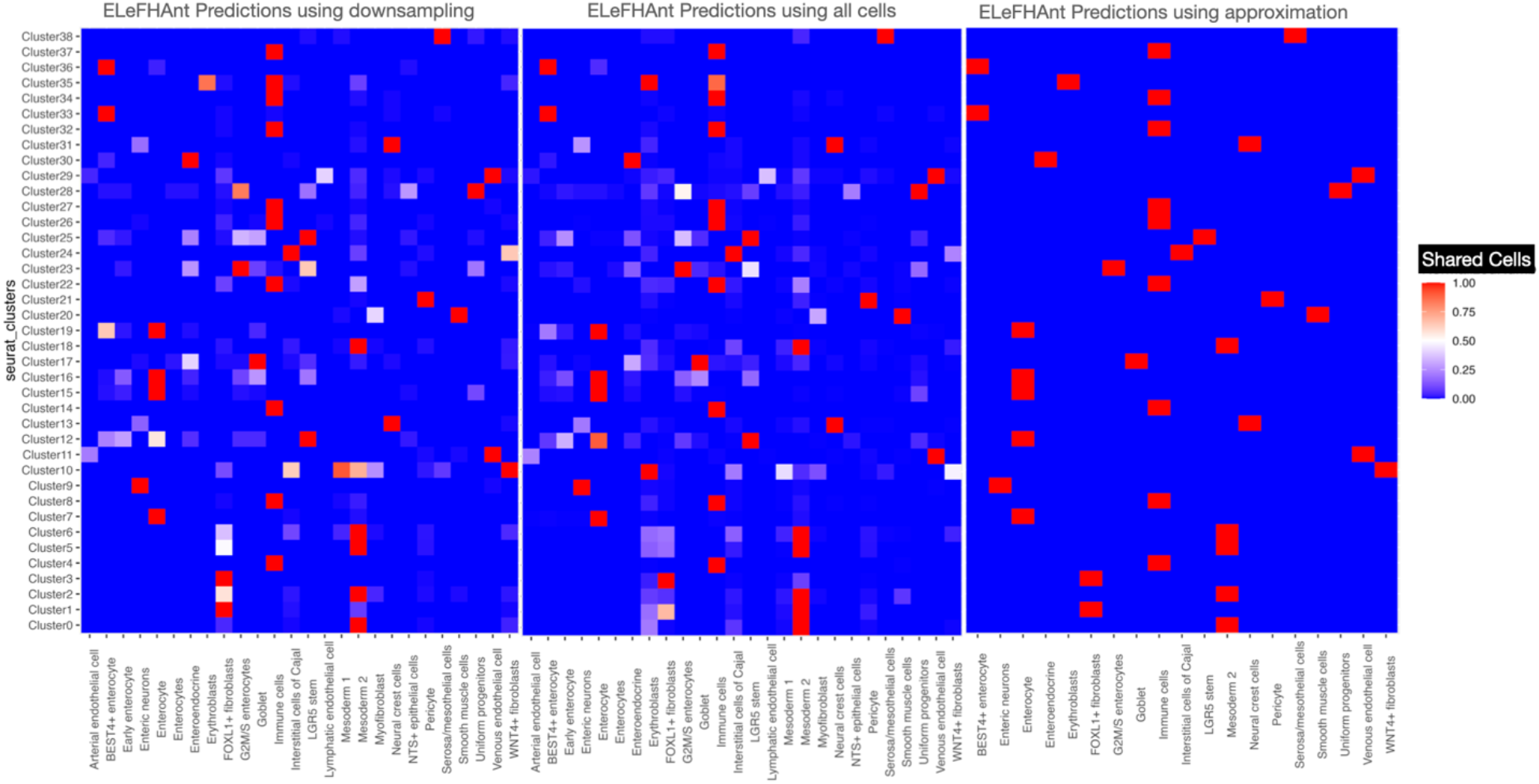
ELeFHAnt choices for each of the scenarios i.e. 1) downsampling reference and query 2) using reference and query unaltered 3) using approximation based prediction

**Supplementary Figure2:**
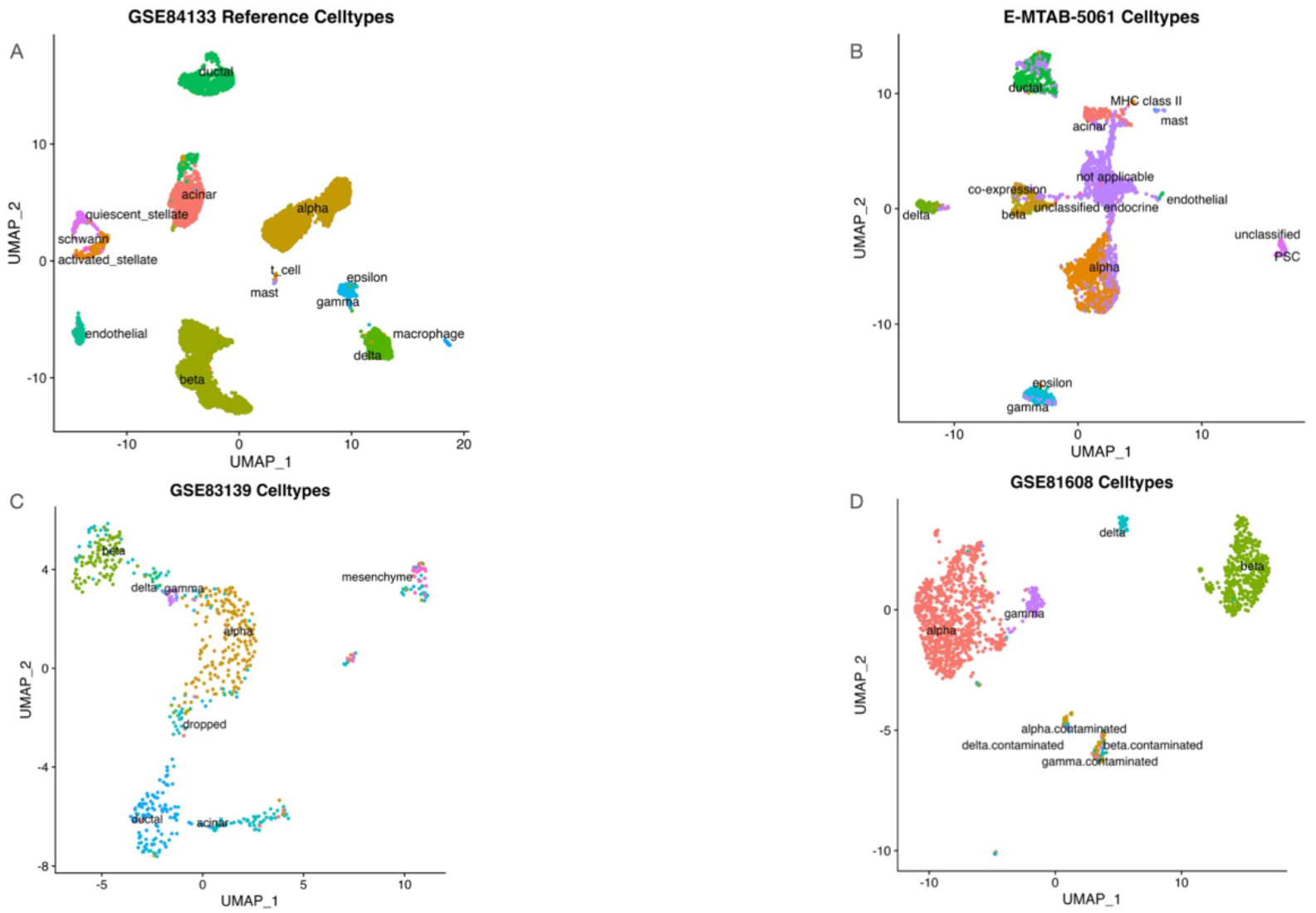
Reference and query cell types in Pancreatic datasets

**Supplementary Figure3:**
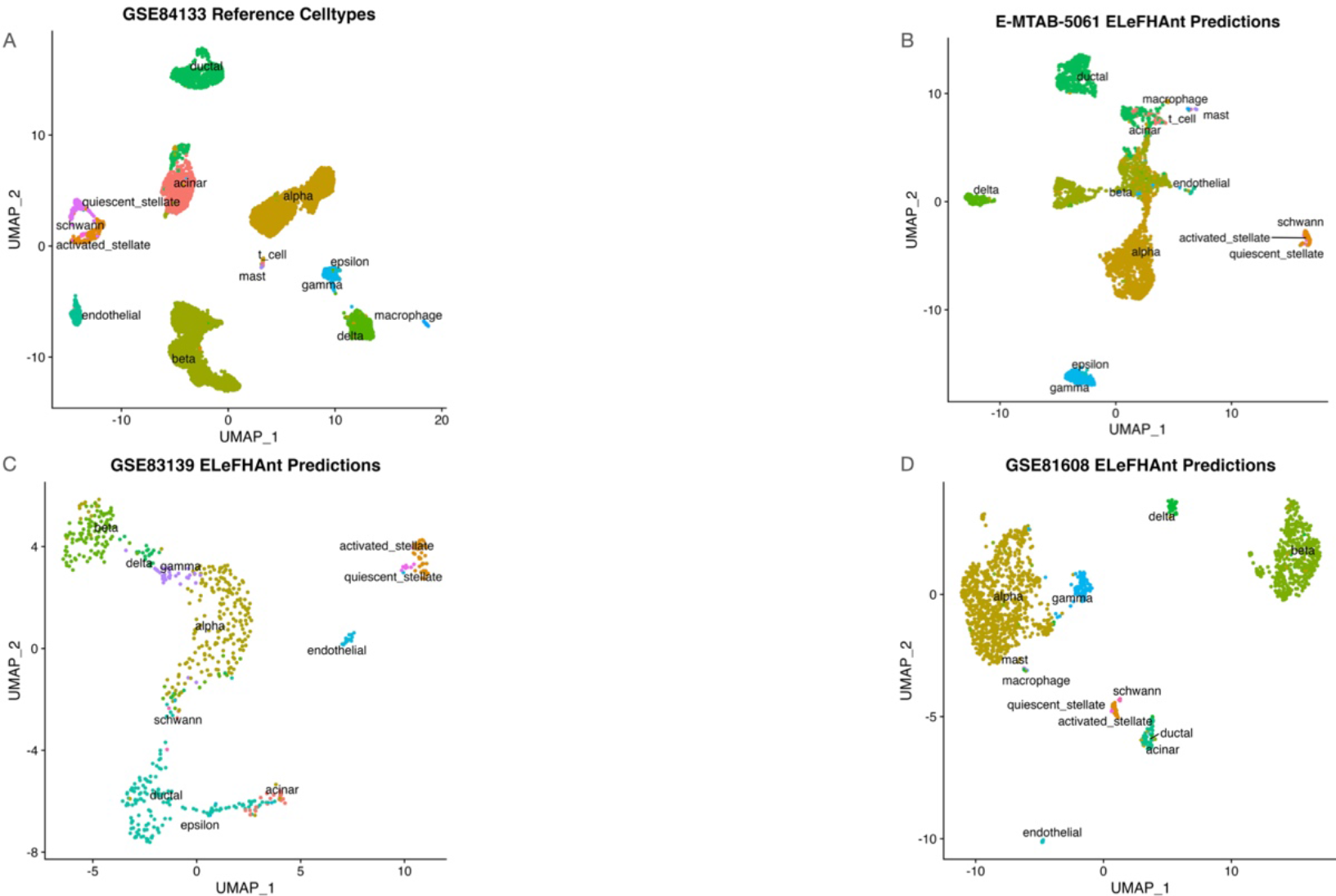
ELeFHAnt predictions for pancreatic datasets

**Supplementary Figure4A:**
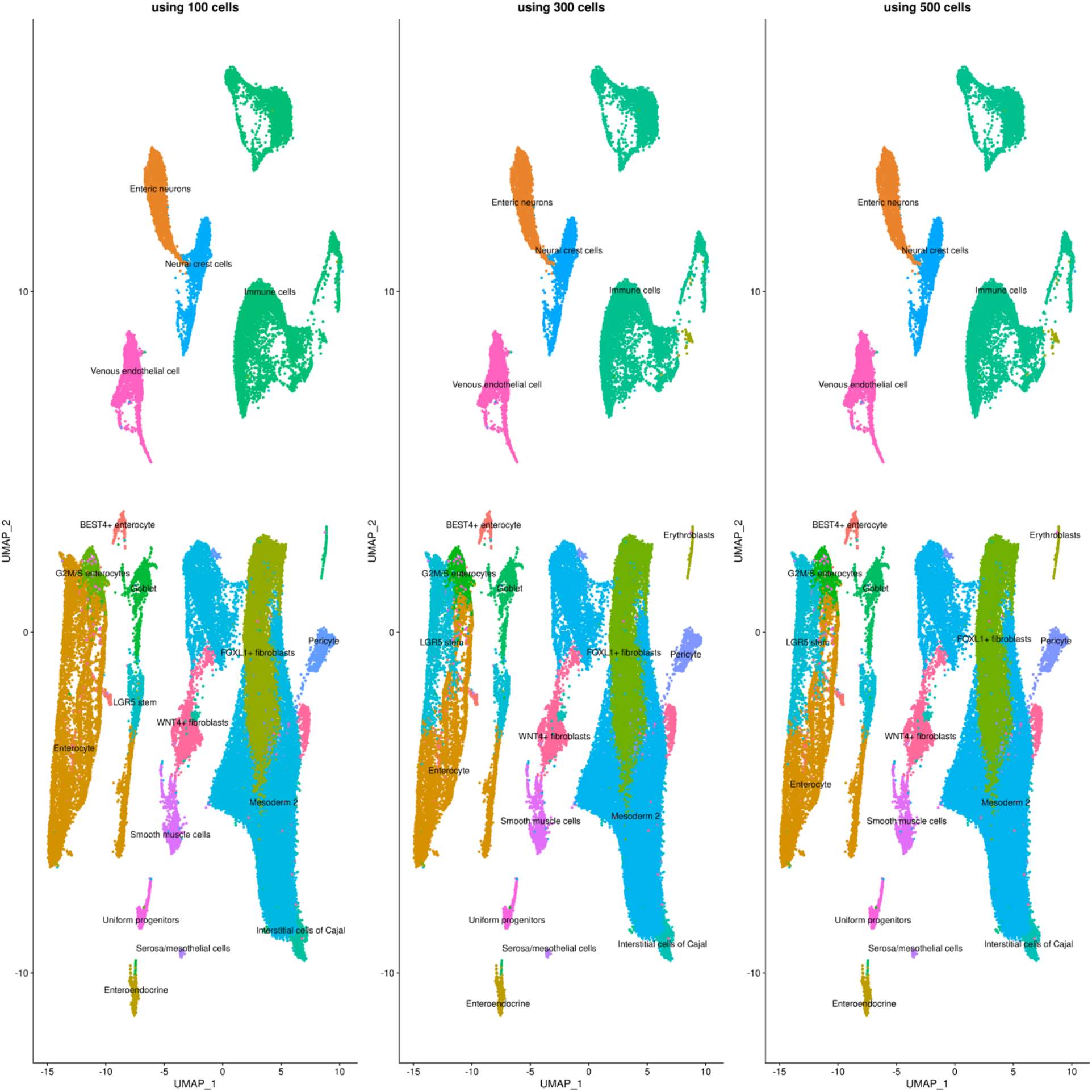
reproducibility test of cell type annotation; subsampling 100, 300 and 500 cells per cell type in the reference and query

**Supplementary Figure4B:**
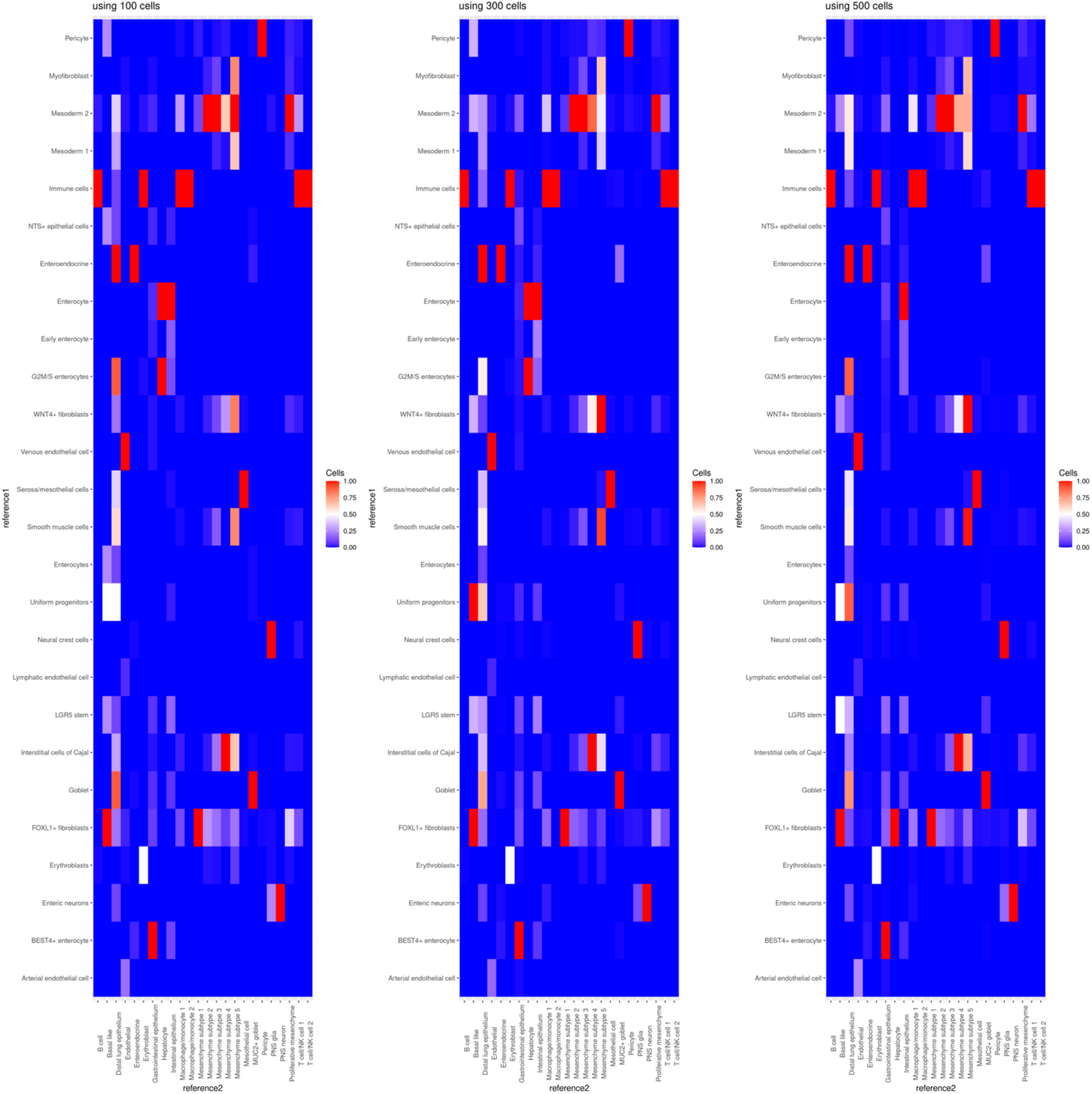
reproducibility test of deduce relationship; subsampling 100, 300 and 500 cells per cell type in the reference1 and reference2

**Supplementary Figure4C:**
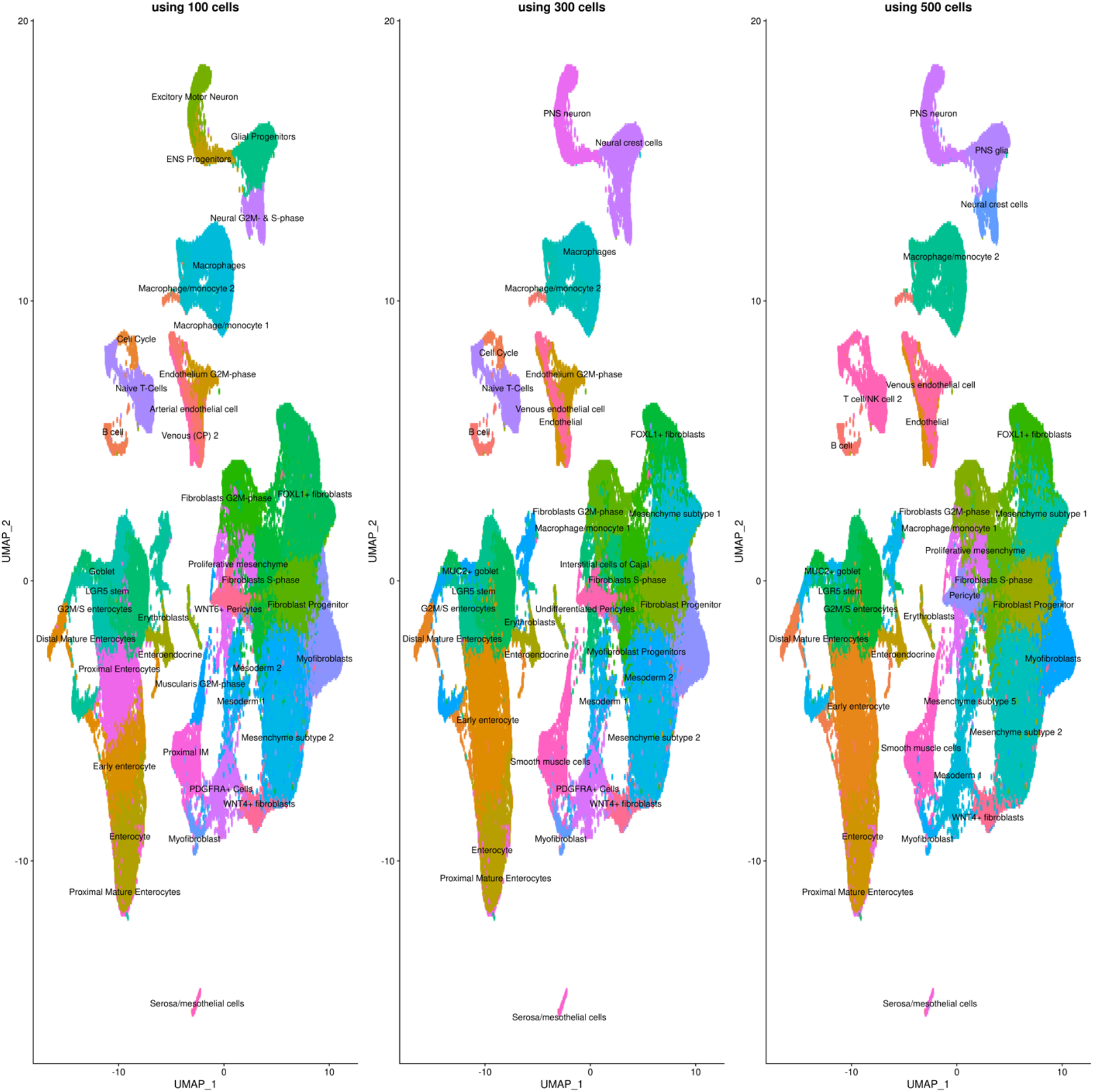
reproducibility test of label harmonization; subsampling 100, 300 and 500 cells per cell label

**Supplementary Figure5:**
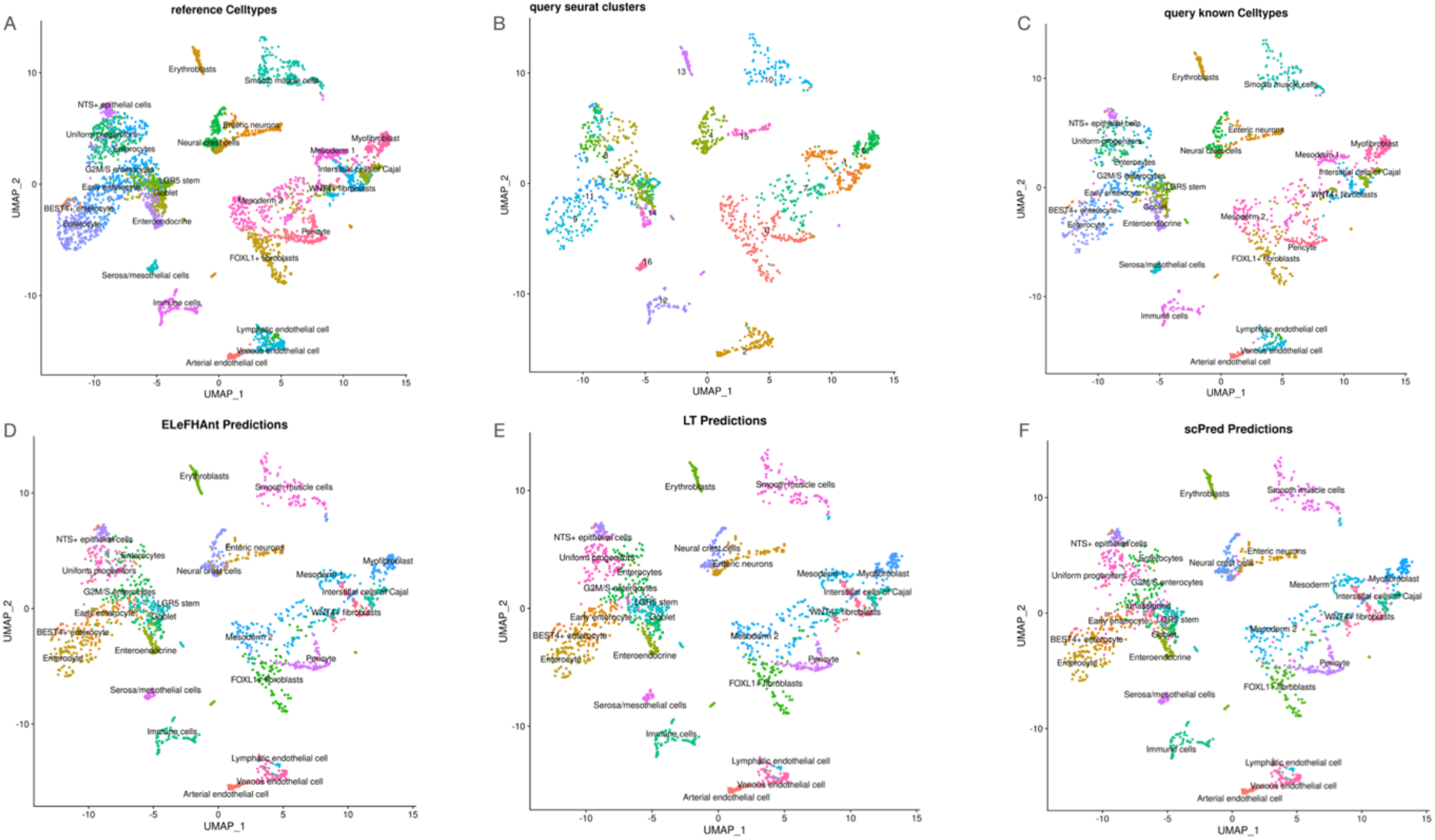
Benchmarking ELeFHAnt, scPred, and Label transfer using Intra dataset training and prediction

**Supplementary Figure6:**
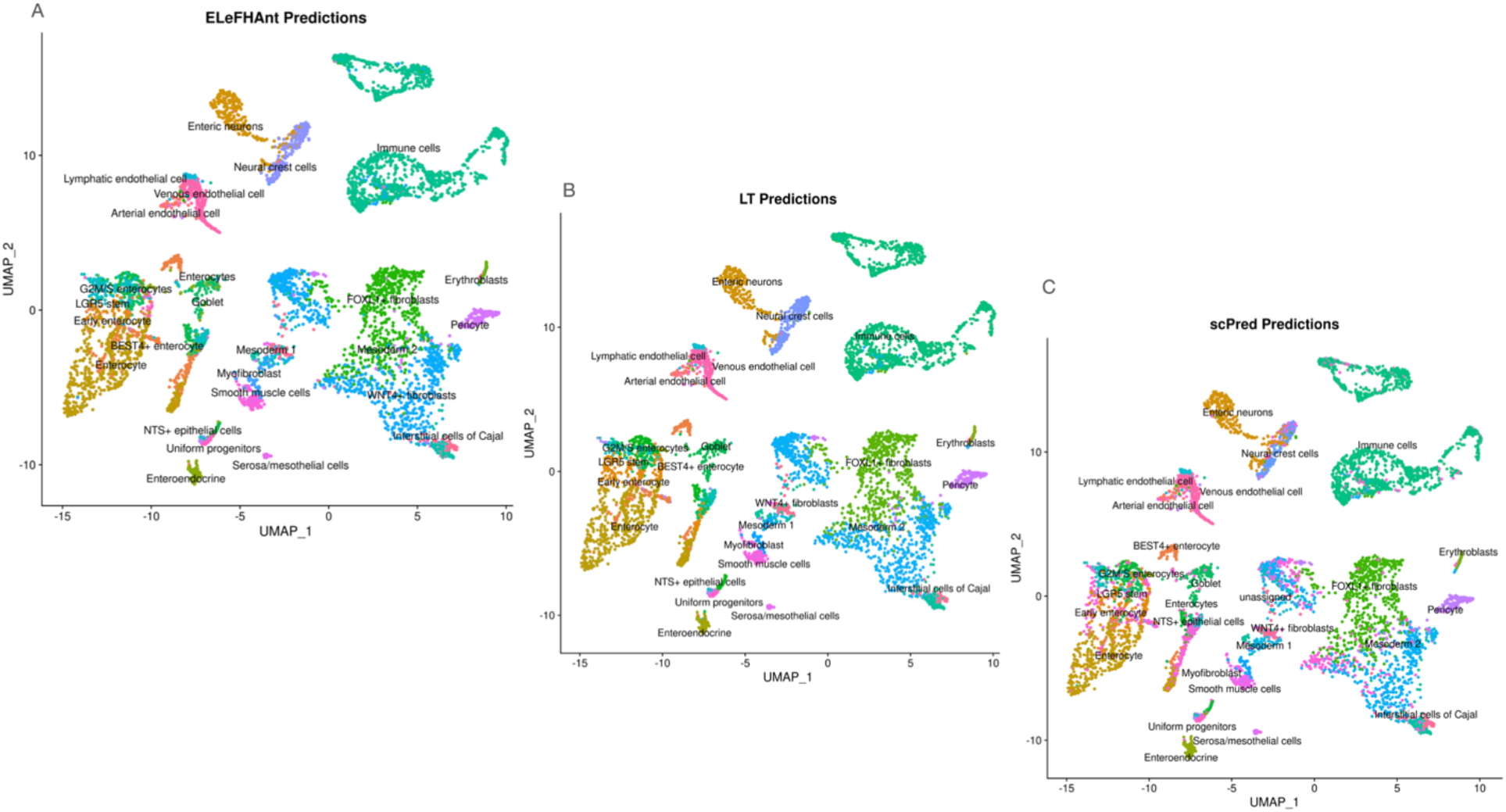
Benchmarking ELeFHAnt, scPred, and Label transfer using Inter dataset training and prediction

**Supplementary Figure7:**
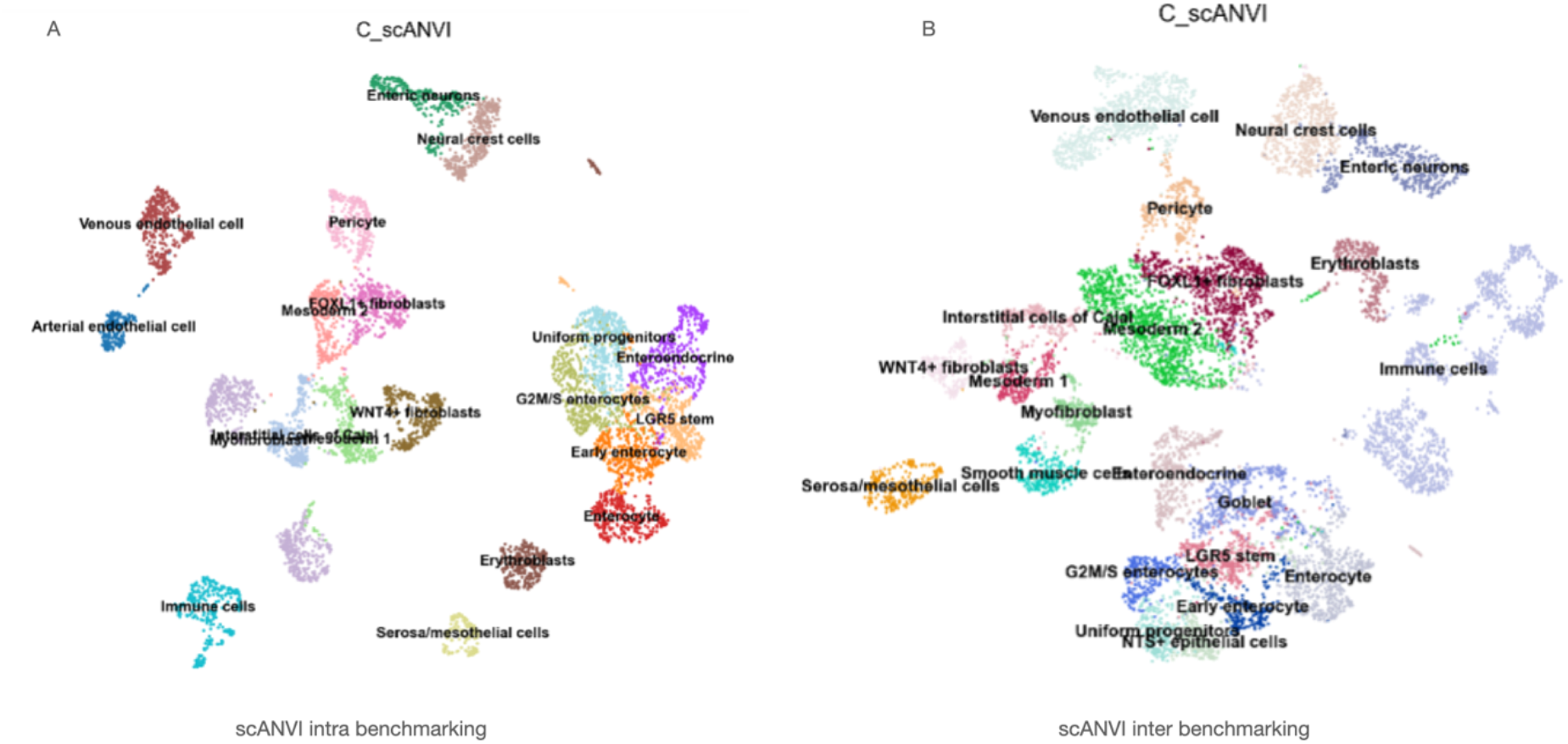
scANVI cell type predictions. A) based on intra dataset training and prediction B) based on inter dataset training and prediction

## References

1. Stuart, T. et al. Comprehensive integration of single-cell data. Cell 177, (2019).

2. The Tabula Muris Consortium., Overall coordination., Logistical coordination. et al. Single-cell transcriptomics of 20 mouse organs creates a *Tabula Muris*. Nature 562, 367–372 (2018).

3. Han, X., Zhou, Z., Fei, L. et al. Construction of a human cell landscape at single-cell level. Nature 581, 303–309 (2020).

4. Pasquini, G., Rojo Arias, J. E., Schäfer, P. & Busskamp, V. Automated methods for cell type annotation on scrna-seq data. Computational and Structural Biotechnology Journal 19, 961–969 (2021).

5. Xu, C. et al. Probabilistic harmonization and annotation of single‐cell transcriptomics data with deep generative models. Molecular Systems Biology 17, (2021).

6. Abdelaal, T. et al. A comparison of automatic cell identification methods for single-cell RNA sequencing data. Genome Biology 20, (2019).

7. Elmentaite, R. et al. Single-cell sequencing of developing human gut reveals transcriptional links to childhood crohn’s disease. Developmental Cell 55, (2020).

8. Fawkner-Corbett, D. et al. Spatiotemporal analysis of human intestinal development at single-cell resolution. Cell 184, (2021).

9. Yu, Q. et al. Charting human development using a multi-endodermal organ atlas and organoid models. Cell 184, (2021).

10. Baron, M. et al. A Single-Cell Transcriptomic Map of the Human and Mouse Pancreas Reveals Inter- and Intra-cell Population Structure. Cell Syst 3, 346–360.e4 (2016)

11. Segerstolpe, Å. et al. Single-Cell Transcriptome Profiling of Human Pancreatic Islets in Health and Type 2 Diabetes. Cell Metab. 24, 593–607 (2016)

12. Wang, Y. J. et al. Single-Cell Transcriptomics of the Human Endocrine Pancreas. Diabetes 65, 3028–3038 (2016)

13. Xin, Y. et al. RNA Sequencing of Single Human Islet Cells Reveals Type 2 Diabetes Genes. Cell Metab. 24, 608–615 (2016)

14. Alquicira-Hernandez, J., Sathe, A., Ji, H. P., Nguyen, Q. & Powell, J. E. scPred: accurate supervised method for cell-type classification from single-cell RNA-seq data. Genome Biology 20, (2019).

15. Meyer, D., Dimitriadou, E., Hornik, K., Weingessel, A., & Leisch, F. e1071: Misc Functions of the Department of Statistics, Probability Theory Group (Formerly: E1071),TU Wien. R package version 1.7-7. (2021).

16. Liaw, A & Wiener, M. Classification and Regression by randomForest. R News 2(3), 18–22, (2001).

